# *Basidiobolus haptosporus-like* fungus as a causal agent of gastrointestinal basidiobolomycosis and its link to the common house gecko (*Hemidactylus frenatus*) as a potential risk factor

**DOI:** 10.1101/311936

**Authors:** Ali Al Bshabshe, Martin R.P. Joseph, Ahmed M. Al Hakami, Tarig Al Azraqi, Sulieman Al Humayed, Mohamed E. Hamid

## Abstract

*Basidiobolus* spp. are a significant causal agent of infections in man and animals including gastrointestinal basidiobolomycosis (GIB). Little information is available on how these infections are acquired or transmitted, apart from the postulation that environmental sources are implicated. This study aimed to identify *Basidiobolus* spp. from GIB patients and from the house gecko as a possible source of infection in Aseer, Saudi Arabia. *Basidiobolus* spp. were isolated from patient specimens (colonic mass biopsy) and from house gecko (gut contents) from Muhayil Aseer areas, in southern Saudi Arabia, using Sabouraud dextrose agar (SDA) which was incubated aerobically for up to three weeks at 30°C. Isolated fungi were initially identified using classical mycological tools and confirmed by sequence analysis of the large subunit ribosomal RNA gene. Cultured specimens from humans and geckos revealed phenotypically similar zygomycete-like fungi which conform to those of *Basidiobolus* species. The strains formed a monophyletic clade in the 28S ribosomal RNA gene phylogenetic tree. They shared 99.97% similarity with *B. haptosporus* and 99.97% with *B. haptosporus* var. *minor* but have a relatively remote similarity to *B. ranarum* (99.925%). One isolates from a gecko (L3) fall within the sub-clade encompassing *B. haptosporus* strain NRRL28635. The study strongly suggests a new and a serious causal agent of GIB related to *Basidiobolus haptosporus*. The isolation of identical *Basidiobolus haptosporus-like* strains from humans and lizards from one area is an important step towards identifying risk factors for GIB. Research is underway to screen more environmental niches and fully describe the *Basidiobolus* strains.

## INTRODUCTION

Basidiobolomycosis is a rare disease caused by the fungus *Basidiobolus*, member of the class zygomycetes, order *Entomophthorales*. It has been isolated from a wide geographical range all around the world (1–3), including Saudi Arabia (4–8). Usually, basidiobolomycosis is a subcutaneous infection, but its etiologic role in gastrointestinal infections has been increasingly recognized (9–12). Gastrointestinal basidiobolomycosis (GIB) is an emerging health care hazard, especially in children. It needs awareness and consideration of differential diagnosis among patients with abdominal masses and eosinophilia, especially in endemic areas such as southern parts of Saudi Arabia (4, 13). Increased perception of basidiobolomycosis and its basic surgical removal along with expanded treatment with itraconazole provide a good possibility for medicinal treatment of the disease (4).

*B. haptosporus* is frequently associated with the gamasid mite *Leptogamasus obesus* (14). Human cases of entomophthoromycosis caused by *B. haptosporus* associate with surgical wounds (15), dermis and subcutaneous tissue (16–20), and invasive mycosis (21). However, one case of gastrointestinal entomophthoromycosis caused by *B. haptosporus* has been documented (14). The authors assumed that many cases of gastrointestinal zygomycosis caused by *B. haptosporus* are either misdiagnosed or pass undiagnosed.

*Basidiobolus* fungi have been linked to the digestive tracts of many amphibians and reptiles, and can be isolated from soil and litter, but most easily from the intestinal contents of reptiles, amphibians and some warm-blooded animals (22–24). In Florida (USA), the occurrence of the fungus in the digestive tracts of *Bufo terrestris, Bufo quercicus, Hyla femoralis, Hyla cinerea, Hyla gratiosa, Hyla squirella, Osteopilus septentrionalis* and *Rana utricularia* has been documented. Species that dwell in terrestrial habitats (*B. terrestris, B. quercicus*, and *R. utricularia*) were found to harbor *Basidiobolus* spp. more frequently (83, 78, and 91%, respectively) than those in arboreal habitats (*H. cinerea, H. squirella, H. femoralis, H. gratiosa*, and *O. septentrionalis* (50, 56, 55, 56, and 70%, respectively) (25). The frequent detection of *B. haptosporus* DNA on or in the gamasid mite *Leptogamasus obesus*, and the absence of the fungus from soil samples seem not to be in line with its assumed ecology as a widespread saprophytic soil fungus. Therefore, a second host species in the life cycle of *B. haptosporus* is discussed as a working hypothesis (26).

Gecko (*Hemidactylus frenatus*) is a reptile or lizard belonging to the family *Gekkonidae* classified in the infra-order *Gekkota*. It is a native species of Southeast Asia and is commonly found in many Asian countries including Saudi Arabia and is called the Asian House Gecko, or simply, House Lizard. There are more than 1,650 species of geckos observed all over the world, especially in tropical warm environments. Geckos (0.64 to 24 inches) are among the most colorful lizards in the world, and appear in various patterns and are easily seen in close association with human habitats in many rural societies around the world (27). A number of research papers have indicated that the house gecko plays a considerable role in the epidemiology of salmonellosis and has implication in public health (28). Nevertheless, there is no available information on its role in transmitting or acting as a source for fungal infections such as basidiobolomycosis.

This study attempts to detect *Basidiobolus* species from an endemic area in Aseer in comparison to a control area. The effort is to understand and determine some of the risk factors associated with this infection.

## MATERIAL AND METHODS

### Sites and collection of samples

The present study included materials ((colonic mass biopsy) from eight human cases clinically suspected of having GIB and gut contents from six captured gecko lizards from Muhayil Aseer (Asir) areas, south Saudi Arabia. Muhayil is situated at 489 m (1832’47.040”N and 422’56.040”E). The climate in Muhayil province is warm in the summer (35-45 °C) and moderate in winter (18-28 °C). Annual rainfall in the province is 30 mm with humidity up to 18%. Muhayil is characterized by low coastal land surrounded by volcanic mountains and valleys with water flow with a natural vegetation cover, which is one of the richest areas of Tehama dense trees (29).

### Diagnosis of gastrointestinal basidiobolomycosis

Eight patients suffering from gastrointestinal basidiobolomycosis (GIB) were included in the study. These patients were admitted between January 2010 and January 2018 to Aseer Central Hospital, Abha, Saudi Arabia. The diagnosis of GIB was made on the basis of clinical and pathological grounds and all have been culture proven. The presented cases had shared characteristics: prolonged fever and symptoms suggestive of chronic infection or malignancies. They were subjected to histopathological and mycological examination. Histological criteria such as granulomatous reaction, dense infiltrate of eosinophils and mycological evidence of fungal structures are taken as suggestive of GIB (30).

Geckos (*Hemidactylus frenatus*) (Figure 1) were captured from different houses at Muhayil Aseer, in southern Saudi Arabia. A small portion of the intestinal contents was collected from each gecko lizard and placed in sterile containers and immediately transported to the laboratory for processing. A total of 6 gecko lizards were included in the study and their gut contents were cultured and processed.

**Figure 1.**
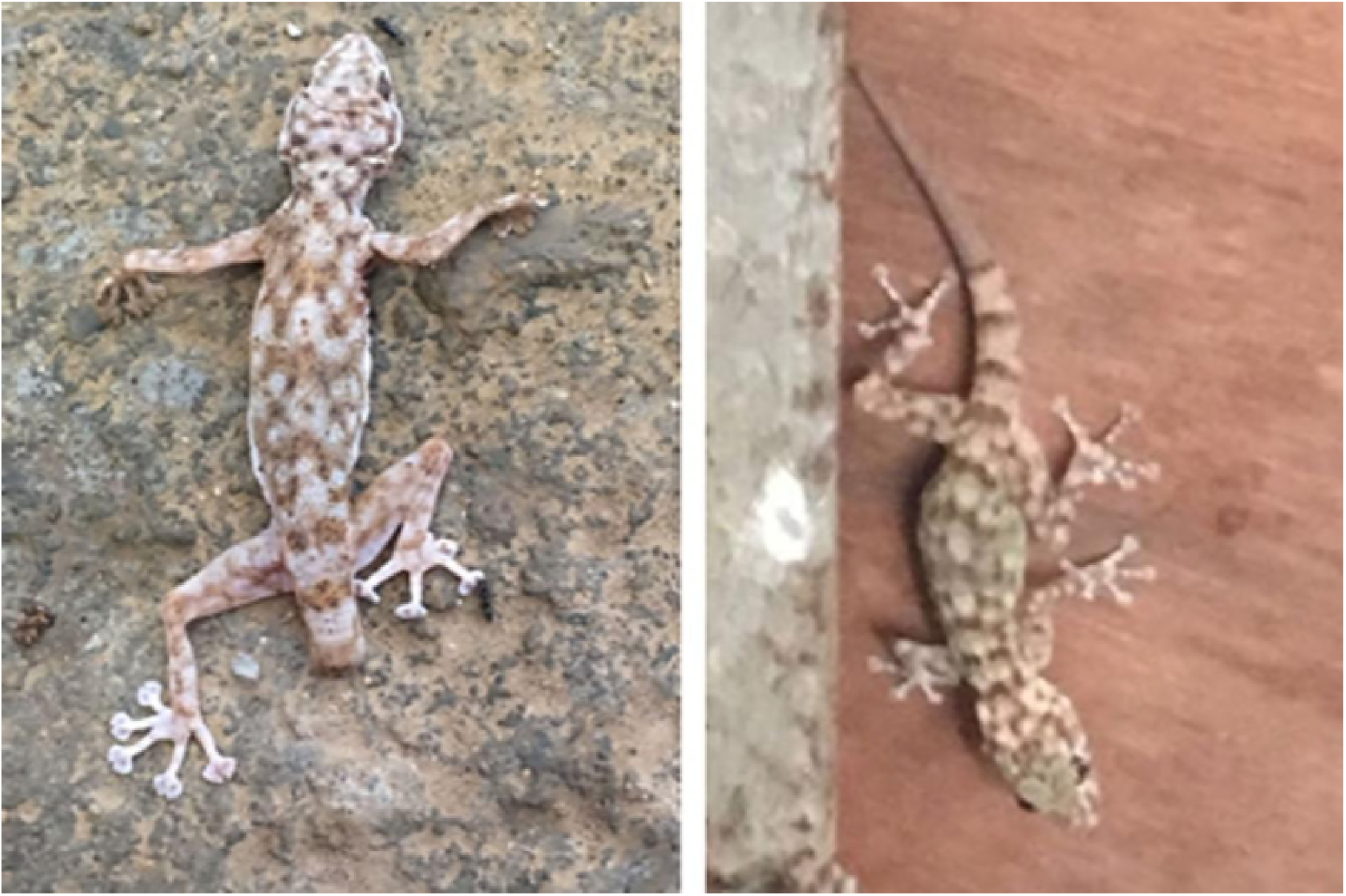
Common house geckos (*Hemidactylus frenatus*) captured from household premises at Muhayil Aseer, Saudi Arabia.

Sabouraud dextrose agar (SDA; Difco Inc.; mycological peptone (enzymatic digest of casein and animal tissues), 10 gm; dextrose, 40 gm; agar, 15 gm; pH adjust to 5.6 at 250 C) was used for original isolation of fungi and for subsequent sub-culturing. Inoculated plates were then incubated at 30°C for up to three weeks. The isolates recovered from infected tissues were examined macroscopically and microscopically. A small piece of grown colonies (thickness, 2 mm; diameter, 2 mm) was placed on lactophenol cotton blue (2 ml phenol, 2 ml lactic acid, 4 ml glycerol, 2 ml H2O) on a clean microscopic slide and examined microscopically.

### Identification of *Basidiobolus* species

Identification of *Basidiobolus* spp. was based on the key of O’Donnell (1) with the following morphological characters as primary for identification of the genus: production of zygospores with or without smooth walls retaining short paired protuberances known as “beaks”, and apical globose primary conidia that were forcefully discharged from the conidiophores, usually still connected to parts of the conidiophore commonly referred to as “skirts”.

### PCR amplification and sequencing of large subunit ribosomal RNA region

DNA amplification and sequencing service was done by Macrogen Inc. (Seoul, Korea). Briefly, the primers LR0R 5′ (ACCCGCTGAACTTAAGC) 3′ and LR7 5′ (TACTACCACCAAGATCT) 3′ (31)were used for amplification of the partial large subunit ribosomal RNA (LSU) region. The PCR reaction was performed with 20 ng of genomic DNA as the template in a 30uLreaction mixture by using an EFTaq (Sol Gent, Korea) as follows: activation of Taq polymerase at 95 °C for 2minutes, 35 cycles of 95 °C for 1minute, 55°C and 72 °C for1minute each were performed, finishing with a 10-minute step at 72 °C. The amplification products were purified with a multiscreen filter plate (Millipore Corp., Bedford, MA, USA). Sequencing reaction was performed using a PRISM BigDye Terminator v3.1 Cycle sequencing Kit.The DNA samples containing the extension products were added to Hi-Diformamide (Applied Biosystems, Foster City, CA). The mixture was incubated at 95 °C for 5 min, followed by 5 min on ice and then analyzed by ABI Prism 3730XL DNA analyzer (Applied Biosystems, Foster City, CA).

### Phylogenetic analysis

The DNA sequences of the strains evaluated were aligned with other reference fungal sequences available in the database using BLAST website, then alignments were inspected visually. The gaps generated were treated as missing data. The fungal DNA sequences were analyzed phylogenetically by the neighbor-joining method using MEGA software (32). Verification for internal branches was calculated by using 100 bootstrapped data sets.

## RESULTS

### Histopathological characterization of GIB tissue samples

Figure. 2 shows a section from a GIB suspected patient. Tissue sections show characteristic histopathology of basidiobolomycosis, including granulomatous reaction incorporating a combination of inflammatory cells, eosinophils and giant cells. The section also shows fungal hyphae and fungal structures surrounded by pink granular materials called Splendore–Hoeppli phenomenon. These criteria are commonly used and enough to confirm clinical diagnosis of GIB.

**Figure 2.**
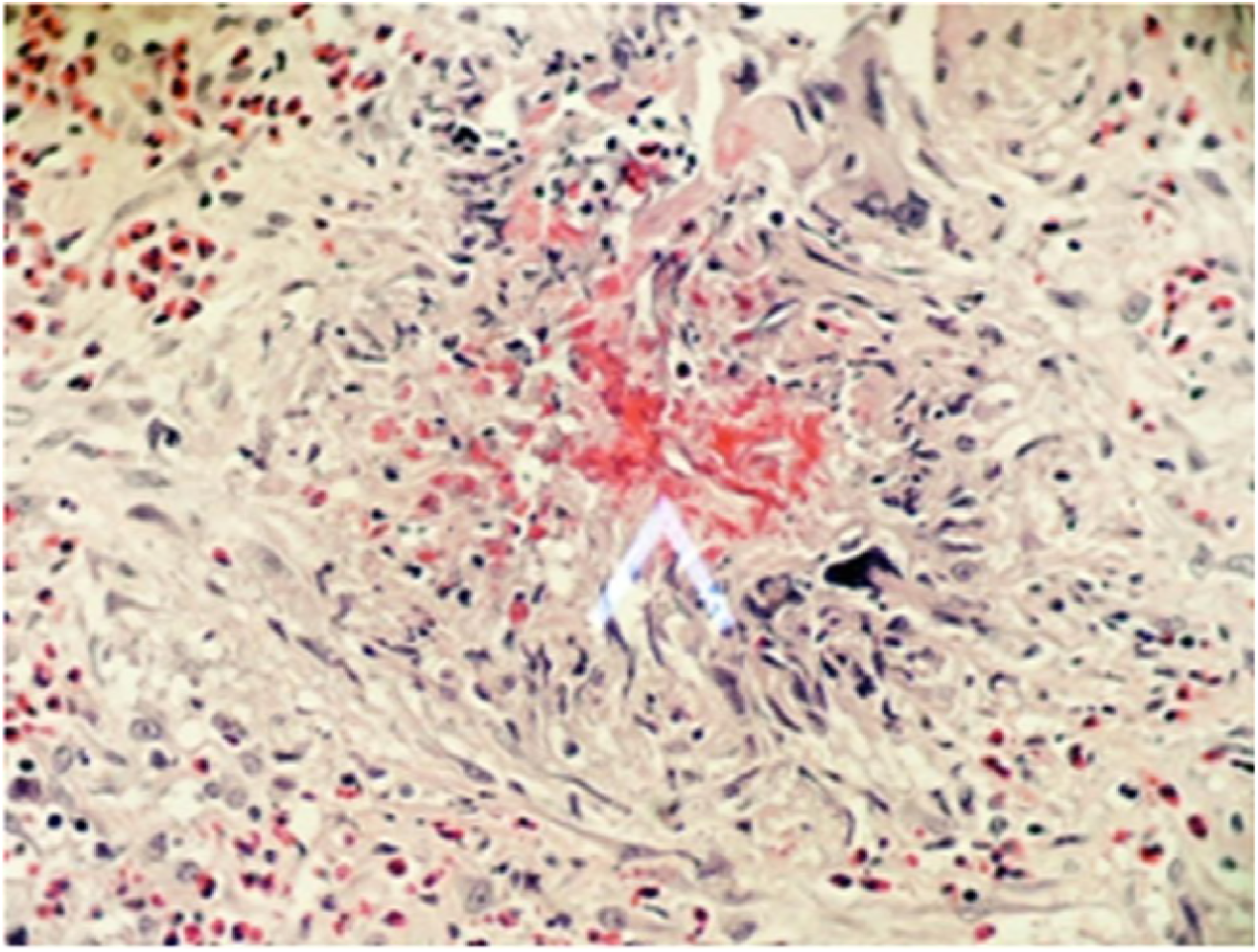
Tissue section from biopsy samples of patient (sample Doza) with GIB showing characteristics of basidiobolomycosis with granulomatous reaction incorporating a combination of inflammatory cells, eosinophils and giant cells. The section also shows fungal hyphae and fungal structures surrounded by pink granular materials called Splendore–Hoeppli phenomenon (white arrow)(H&E stain).

### Morphological identification of *Basidiobolus* species

Eight isolates were recovered from suspected GIB cases and 6 from gut contents of the common house gecko (*Hemidactylus frenatus*). The phenotypic characterization of the 14 isolates was completed. Culture proven human isolates were: ARD (2010); BoTi (2011); Doza (2013); 9-4 (2014); V81 (2017); ACH (2017); F43-5 (2016); F15-1 (2017) and the gecko’s culture proven isolates were: L1 (2018); L3 (2018); L4 (2018); L4G (2018); L6 (2018) and L7 (2018).

The organisms developed fast-growing isotropic, pale colonies in the primary culture on SDA at30°C (Fig. 3). On this medium, within the first 3 days of incubation at either temperature, flat membranous colonies with a smooth, glabrous, and waxy appearance developed. Older colonies (3 days old and older) became powdery in appearance, with short aerial mycelia, and developed radiate folds from the centers of the colonies (Fig. 3). Morphological characteristics conform to *Basidiobolus* species that were round, flat, waxy, glabrous and radially folded colonies (33, 34). The tested strains were found to have phenotypic properties distinctive for members of the genus *Basidiobolus*. The 12 isolates from human cases of GIB and from geckos were identified as *Basidiobolus* species based on their macroscopic and microscopic features.

**Figure 3.**
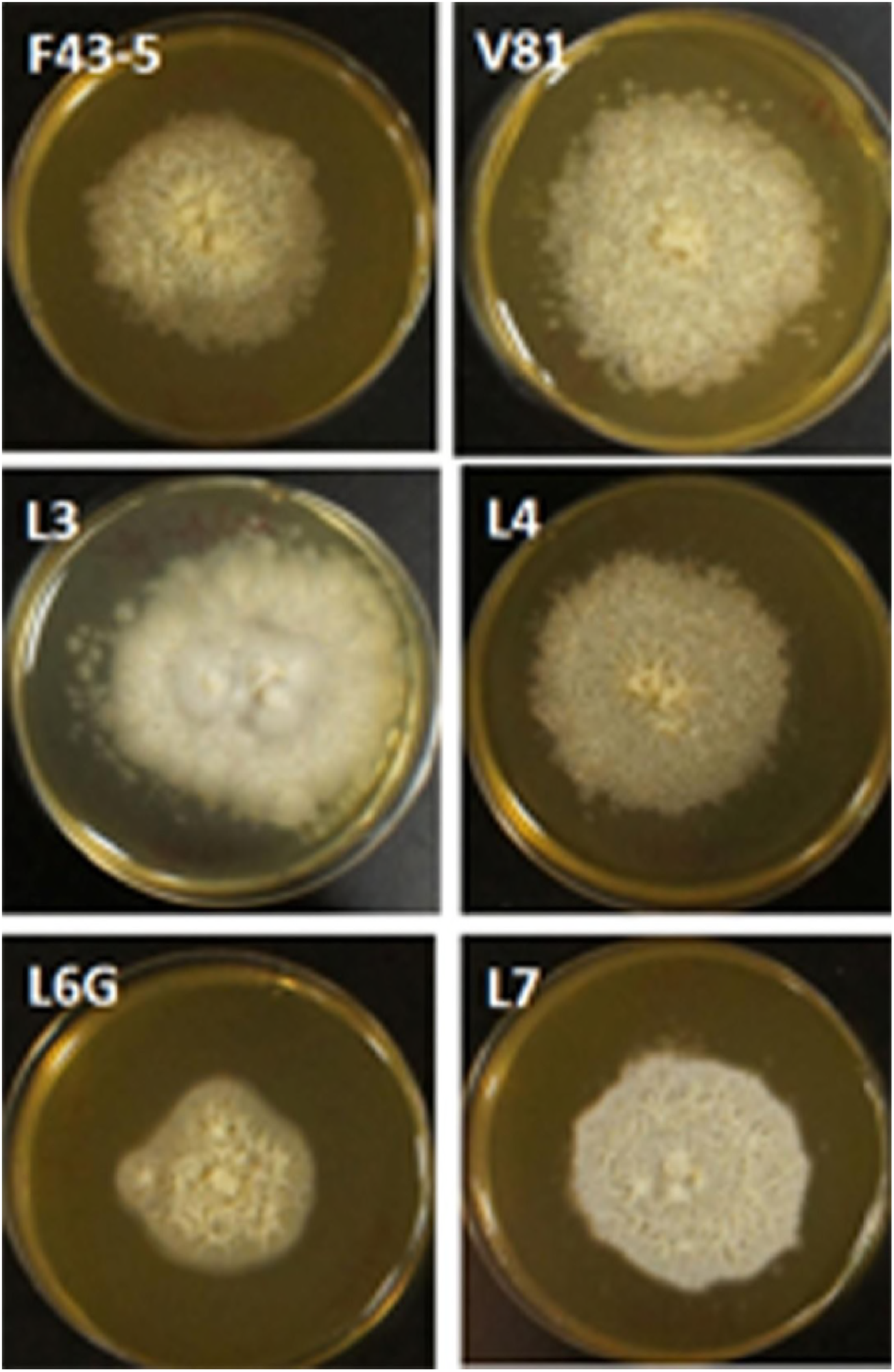
Growth of a 5-day old culture of *Basidiobolus* spp. on Sabouraud’s dextrose agar at 30°C showing yeast-like colonies, smooth, glabrous in texture and creamy yellow.

Microscopically, young colonies showed colorless broad hyphae with few septa, with smooth, thick walls, and abundant large, spherical, darkly colored chlamydospores and zygospores are formed. Microscopically, colonies were examined from 3 days and 5 days SDA growth. They showed hyaline, coenocytic, ribbon-type, unbranched hyphae measuring 5.0 to 10.0 (2 to 4) μm in diameter, with occasional septa, usually on the distal parts of the hyphae (Fig. 4). Immature zygospores developed after the encounter of two hyphal segments, producing swelling in the contact area and later becoming globose, with several internal vacuoles (data not shown). Numerous smooth globose to subglobose ripe zygospores, 10.0 to 18.0 um in diameters, were observed on primary SDA cultures for some strain, but not for the other two strains. The internal cell walls of the mature zygospores were 1.0 to 4.0 um in diameter (0.5 to 1.0 um; n _ 50). Some of the mature zygospores contained a readily detectable large globule at the center, with some space between the internal globule and the cell wall (Fig. 4).

**Figure 4.**
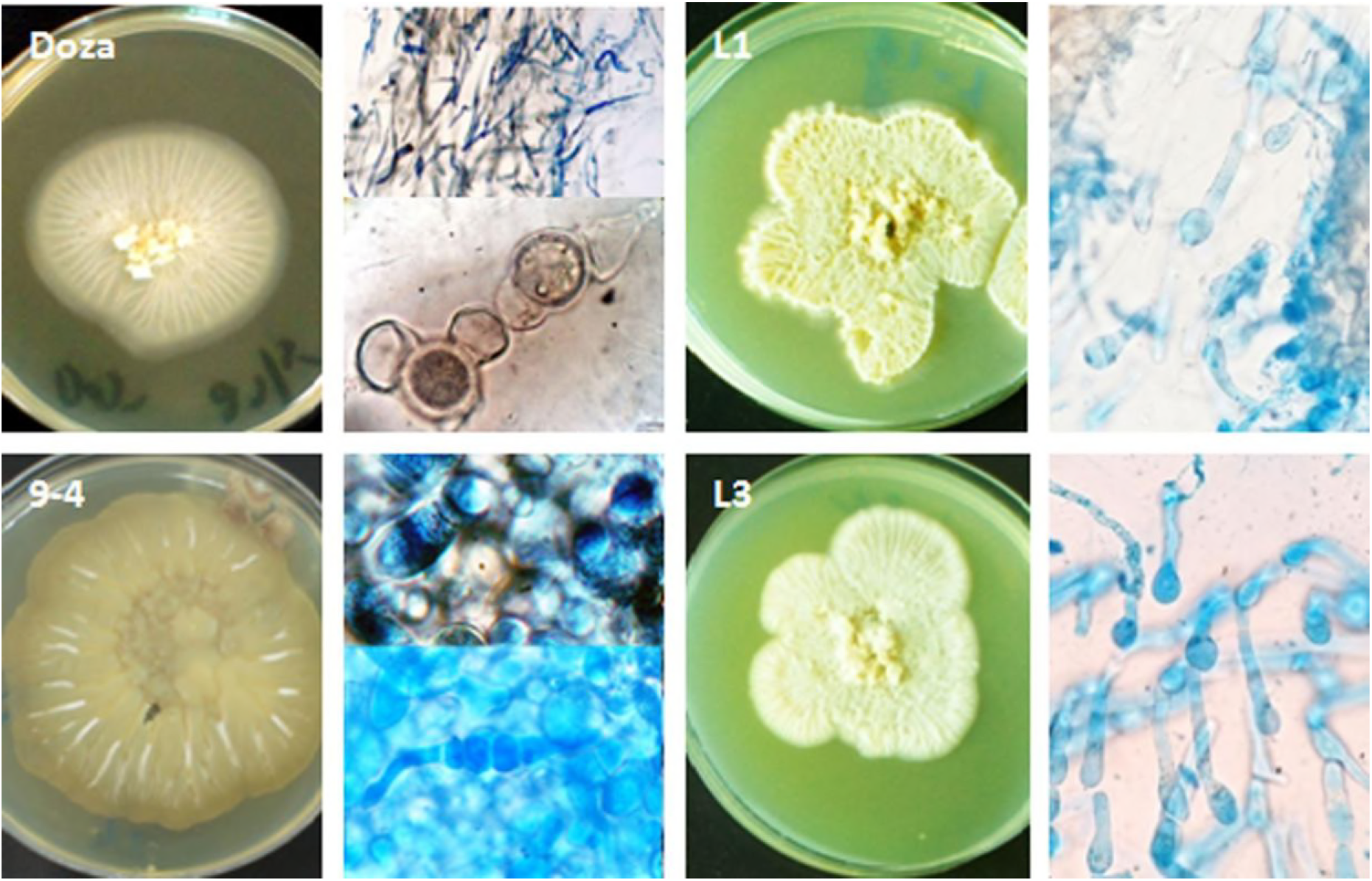
Colony morphology and microscopic appearance of some *Basidiobolus* strains. Note the broad hyaline, coenocytic hyphae characteristic of *Zygomycetes*, with numerous smooth, globose mature zygospores stained by lactophenol cotton blue.

### Phylogenetic analysis of *Basidiobolus* species

Comparison of the 28S rDNA sequences of the isolates with corresponding nucleotide sequences of representatives of the order *Entomophthorales* (35) confirmed that they belong to the genus *Basidiobolus*. Isolates Doza, 9-4, V81, F43-5, F15-1, L1, L4 and L4G were found to have identical 28S rDNA sequences. The strains formed a monophyletic clade in the 28S ribosomal RNA gene. They shared 99.971% similarity with *B. haptosporus* NRRL28635 and shared 99.969% with *B. haptosporus* var. *minor* strain ATCC 16579 and remotely held similarity to *B. ranarum* (99.925%) (Figure 5). The high 28S rDNA gene sequence similarities to the representatives of the genus *Basidiobolus* (93.9 to 98.7%) showed by these isolates support their addition to this genus. Few nucleotide mismatches were found within the isolates Doza, 9-4, V81, F43-5, F15-1, L1, L4 and L4G DNA sequences from humans and gecko lizards. One isolate from gecko (L3) fall within the sub clade encompassing *B. haptosporus* strain NRRL28635. High-stringency BLAST (http://www.ncbi.nlm.nih.gov/) analysis of the three sequences showed strong identity of the human isolates with those from lizards. *Conidiobolus* species were used as an out-group, and the constructed phylogenetic trees constructed by either neighbor joining, minimum evolution, or parsimony demonstrated no substantial topological difference.

**Figure 5.**
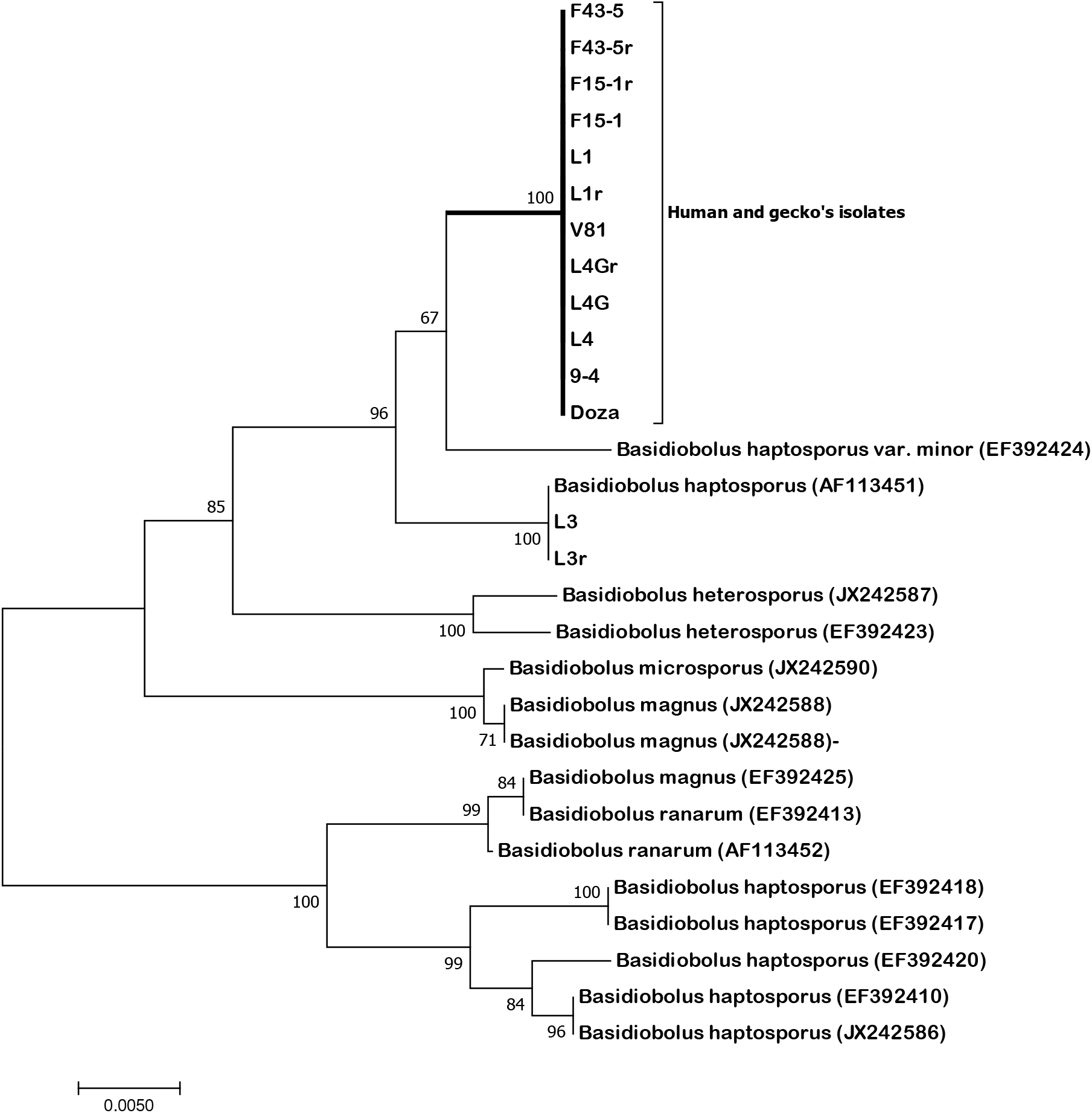
Neighbor-joining tree of aligned 28S large subunit ribosomal RNA genes of the five strains from humans (9-4, Doza, V81, F43-5, F15-1) and 3 strains from gecko lizards (L1, L4, L4G). The percentages of 100 bootstrap re-sampled data obtained by neighbor-joining analysis are given on the branches. Accession numbers of the DNA sequences of our strains and the reference strains retrieved from GenBank are given in brackets. *Basidiobolus haptosporus*-like fungus DNA sequences assembled in a monophyletic group connected to *B. haptosporus* and *B. haptosporus* var. *minor*. The bar represents 0.005 substitutions per nucleotide.

## DISCUSSION

The present study aimed to identify fungal isolates from patients with symptoms and signs simulating GIB and attempted to isolate similar fungus from a potential risk lizard, the house gecko. The assumption that house geckos could be a contaminating agent for people in the rural houses in Aseer region was made after a lot of considerations. The validity of the question is based on two facts; 1) almost all cases of GIB originated from Muhayil province and that Abha region a neighboring high altitude region reported no or one case of GIB; 2) gecko population especially in household premises is not usually seen in Abha, contrary to Muhayil where household premises contain many geckos seen almost in every house hold.

The tested strains recovered from humans and geckos were have phenotypic properties distinctive for members of the genus *Basidiobolus* (33, 34). The eight isolates from human cases of GIB studied over the last 8 years were identified as *Basidiobolus* species based on their macroscopic and microscopic features. Our initial phenotypic comparison of *Basidiobolus* spp. with that isolated from house gecko (gut contents) did not reveal a solid conclusion on their close similarity, since phenoltyping alone is always not conclusive at specie level (33). Application of DNA sequence analysis, for instance, using 28S rDNA gene was found very useful. This was useful in the present study supporting our hypothesis, as human and lizard isolates clustered in one clade related to but were readily distinguishable from *B. haptosporus* and *B. haptosporus* var. *minor* (Fig. 5). The results show that *Basidiobolus* isolates from humans and geckos from Aseer region have identical partial 28S rDNA sequences that distinguish them from representatives of closely related taxa notably *B. haptosporus* and *B. haptosporus* var. *minor*.

Basidiobolomycosis mostly manifests as subcutaneous infections, but progressively increased gastrointestinal and other systemic infections have been observed (9–12, 30). Gastrointestinal basidiobolomycosis (GIB) is an emerging health care hazard, especially among children in tropical countries. The role of *Basidiobolus ranarum* in GIB has long been suspected (3, 22, 36, 37). Diagnosis of the majority of basidiobolomycosis cases is established on histological basis, fungal morphology and the Splendore-Hoeppli phenomenon (30, 34, 38, 39). Not all studies provided enough culture proven cases. This situation prompted the need not only for culture but DNA based confirmation of the type of these zygomycetes (40).

A number of research papers have indicated that the house gecko plays a considerable role in the epidemiology of salmonellosis and has implication in public health (28). Nevertheless, there is no available information on its role in transmitting or acting as a source for fungal infections such as basidiobolomycosis.

*B. haptosporus* is frequently associated with the gamasid mite *Leptogamasus obesus* (14). Human cases of entomophthoromycosis caused by *B. haptosporus* associate with surgical wounds (15), dermis and subcutaneous tissue (16–20), and invasive mycosis (21). However, one case of gastrointestinal entomophthoromycosis caused by *B. haptosporus* has been documented (14). The authors assumed that many gastrointestinal zygomycetes caused by *B. haptosporus* are either misdiagnosed or pass undiagnosed.

### Conclusions

Our work identified *B. haptosporus-like* fungus from all the 3 clinically-diagnosed GIB patients. These isolates are not related to *B. ranarum*, previously linked to this disease. The study identified house geckos as a source of infection given the abundance of the fungus in their guts. This type of lizard is not commonly seen in Abha district, as compared to Muhayil district which has both GIB and a lizard population in almost every house and surrounding premises. This study reports a link between human GIB and gecko lizard as a source of infection and the identification of *B. haptosporus-like* fungus as a cause of human GIB. Study is going on to further screen more lizard samples and their environmental habitat including water resources and soil in the GIB endemic area. Full detailed of these isolates also require detailed analysis.

### Nucleotide sequence accession numbers

The partial 28S rDNA sequences determined in this study for nine strains have been deposited in the GenBank database under accession numbers Strain Doza, MH254938; 9-4, MH256645; L4, MH256646; L4G, MH256647; V81, MH256648; L1, MH256649; F15-1, MH256650; F43-5, MH256651 and L3, MH256652.

